# Single-sequence deep learning delivers crystal-quality models of covalent K-Ras G12 hotspot complexes

**DOI:** 10.1101/2025.09.16.676163

**Authors:** Sungwon Jung, Qinheng Zheng, Kevan M. Shokat

## Abstract

Structure-based design of covalent drugs has achieved tremendous success by understanding and leveraging the three-dimensional interactions between small-molecule drug candidates and their protein targets. However, this approach traditionally relies on high-resolution co-complex structures obtained by X-ray crystallography, NMR, or cryo-EM, methods that are time-consuming and resource-intensive. Here we show that Chai-1, a publicly available structure prediction tool that accepts user-defined ligands, is able to accurately predict covalent K-Ras(G12C) complexes without using a multiple sequence alignment (MSA). Chai-1 yields pocket-aligned RMSDs <2 Å for chemically diverse K-Ras(G12C) inhibitors, ranging from ARS-853 to BBO-8520. In addition to the conventional acrylamide-based covalent K-Ras(G12C) inhibitors, Chai-1 with a covalent-bond restraint successfully reproduced the binding poses of covalent K-Ras(G12D) and K-Ras(G12S) inhibitors, while showing limitations in capturing chemical details such as accounting for leaving-groups, bond properties, and stereochemistry. Chai-1 also provides ∼40-fold higher throughput than state-of-the-art AlphaFold3 while maintaining comparable pose accuracy. Together, these findings establish Chai-1 as an accessible and computationally efficient tool for covalent protein-ligand co-complex structure prediction, with its covalent-restraint mode offering a unique solution to accelerate covalent drug discovery, especially for challenging targets beyond cysteine.

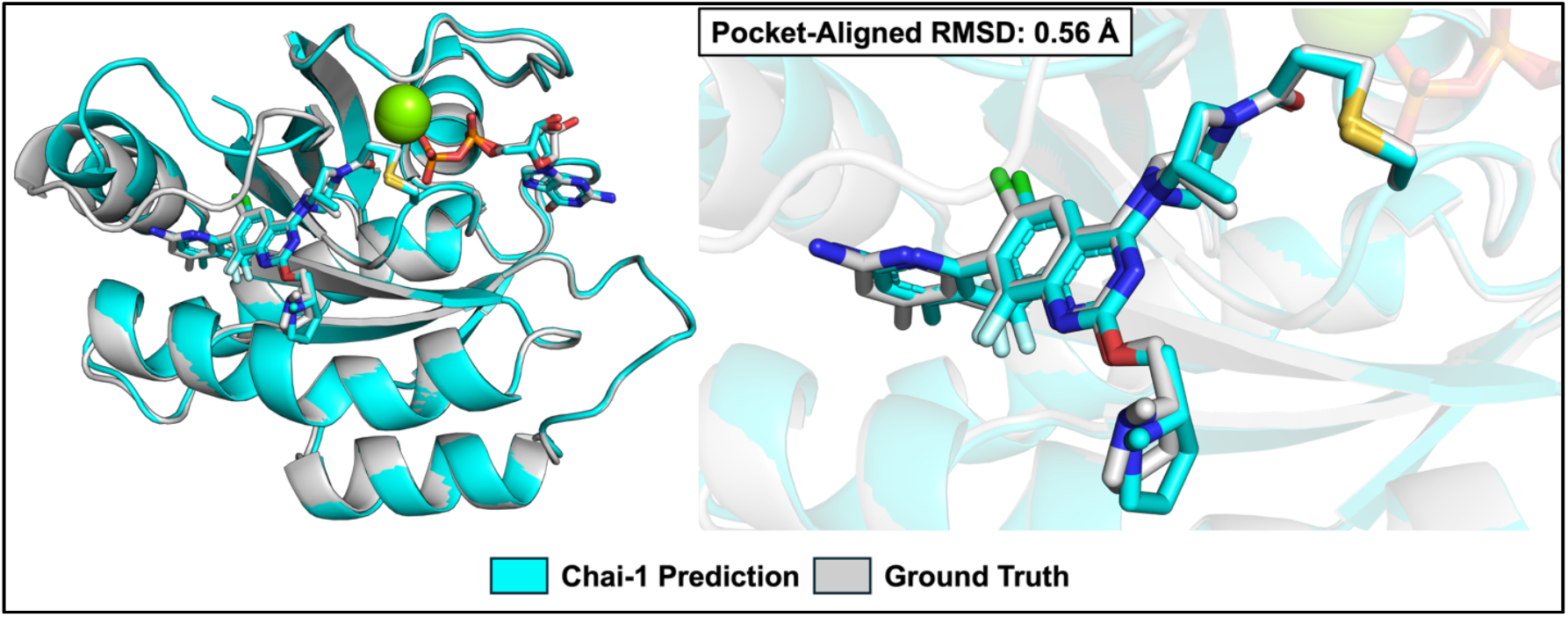

## INTRODUCTION

*KRAS* is the most frequently mutated oncogene that drives ∼30% of human cancers^1^. The *KRAS* oncogene encodes a small GTPase K-Ras, which was historically considered to be an “undruggable” target for cancer therapy because of the lack of small molecule binding sites^2^. Acrylamide-based Compound 12^3^ overturned that view by forming a covalent bond with Cys12 of one of the most common K-Ras missense mutations, K-Ras(G12C), revealing a previously unknown small molecule binding pocket. Now known as the switch-II pocket^3–5^, the space between Switch-II and α3 helix has been exploited by medicinal chemists to deliver an increasing number of drug candidates, including two FDA-approved K-Ras(G12C) covalent inhibitors to treat non-small cell lung cancer, sotorasib and adagrasib.

Despite these breakthroughs, the drug development pipeline still relies on co-crystallography, a step that could lag chemical optimization by weeks or months^6^. This bottleneck is particularly pronounced for dynamic protein targets, presenting a significant hurdle for the rapid design and iteration of novel covalent inhibitors^7^. This rising demand for faster, more efficient structural insights has fueled the development of purely in silico routes, such as all-atom MD^8^ and covalent-docking workflows^9,10^, which could supply plausible binding-mode models. Deep-learning approaches are beginning to demonstrate success alongside traditional physics-based docking methods, often yielding more accurate structural predictions for complex covalent ligand-protein interactions^11–13^.

Deep-learning-based biomolecular complex-structure prediction models like AlphaFold3 (AF3)^12^ could in principle supplement crystallization or spectroscopy-based experimental structural determination. However, the public AlphaFold Server has clear restrictions: it can be used only for non‐commercial purposes and prohibits the use of server outputs in docking or screening tools, while also accepting only a curated list of biologically common ligands rather than arbitrary, user‐defined small molecules. These limitations significantly reduce its applicability in drug‐discovery campaigns. In contrast, Chai-1^14^ specifically fills this gap: its free web server accepts user-specified ligand SMILES along with any protein sequence and returns structure models in mmCIF (.cif) format, with the option to compute them in “single-sequence” mode without requiring multiple-sequence alignment (MSA) or templates. This makes it straightforward to move from sequence and SMILES to docking. In addition to being fully available for commercial use under an Apache 2.0 license, the open-source code supports covalent-bond restraints between the ligand and protein, enabling the use of the covalent bond as a constraint to aid in solving less common electrophile warheads and more challenging covalent amino acid targets such as aspartate and serine.

Our lab has been interested in developing covalent ligands targeting the somatically mutated oncogenic K-Ras variants, including G12C^3^, G12S^15^, G12R^16^, and the most frequent G12D^17,18^ mutation. Our work benefited greatly from an early co-crystal structure of a tethering compound we discovered covalently bound to K-Ras(G12C)^3^, revealing a pocket not visible in previous PDB structures due to the dynamic nature of the Switch II loop. As our work evolved beyond G12C, we were often challenged by a lack of ability to obtain structures for ligands that reacted too slowly or incompletely, making optimization difficult. Once more optimized compounds were obtained, we were able to solve helpful co-crystal structures. The recent explosion in machine learning based structure prediction attracted our attention, and we became curious if these methods could accurately predict structures we had recently solved (including one that had not yet been deposited when we initiated this work; PDB: 9DMM^19^). We thought this exercise would allow us to benchmark current ML methods, and since we are not a computational lab, we hoped our experience might be helpful to similar groups. Because the target pocket remained constant while only the nucleophilic residue varied (Cys, Asp, or Ser), we were able to attribute differences in prediction accuracy to the covalent features.

Here, we show that Chai-1 accurately predicts K-Ras(G12C) co-crystal structures, extends to difficult targets such as K-Ras(G12D) and K-Ras(G12S) using covalent restraints, and achieves superior computational efficiency compared to AlphaFold3 without substantial loss of accuracy. Notably, three of the hotspot mutations at codon 12 (G12C, G12D, and G12S) now have published covalent ligands, providing a controlled test in which the switch-II pocket geometry is well characterized and the primary variable is the electrophile chemistry. By benchmarking across these variants, we asked whether Chai-1 could faithfully reproduce the experimentally determined binding poses for diverse warheads, and in what contexts the predictions deviated. Such a framework will be particularly valuable as the field advances beyond cysteine toward more challenging covalent amino acid targets, where experimental structures remain scarce and computational predictions will likely provide the primary guidance for covalent ligand design.

## RESULTS

### Crystal-Quality Web Predictions for K-Ras(G12C) Switch-II Pocket Inhibitors

We first asked how Chai-1 would perform on our recently determined 1.90-Å K-Ras(G12C)– Divarasib (Figure 1) structure^19^, which had not been available in the PDB at the time of prediction, serving as a true blind test. The Chai-1 web mode delivered an overlay with a pocket-aligned RMSD of 0.56 Å (Figure 2a), surpassing the 2.0 Å-threshold commonly used in deep learning–based docking benchmarks^12^. Consistent with its well-known conformational plasticity^20^, the switch-II loop is the only region where the Chai-1 prediction substantially deviates from the crystal structure. Notably, this segment receives comparatively lower predicted local distance difference test (pLDDT) scores, although it remains within the high-confidence range (Figure 2b). Furthermore, while examining the outputs we noticed that in several restraint-free predictions, PyMOL displayed a covalent link between Cys12 and the acrylamide warhead even though no covalent bond restraint had been applied. Follow-up checks confirmed that this bond was not generated by Chai-1 itself, but was instead drawn by PyMOL’s default heuristic, which infers connectivity from inter-atomic distances and element types when explicit CONECT records are missing in the input file. We nonetheless note that the appearance of such “spontaneous” bonds can be taken as a fortuitous hint of a strong intrinsic preference for covalent engagement and could serve as a qualitative cue when prioritizing covalent drug candidates.

**Figure 1.**
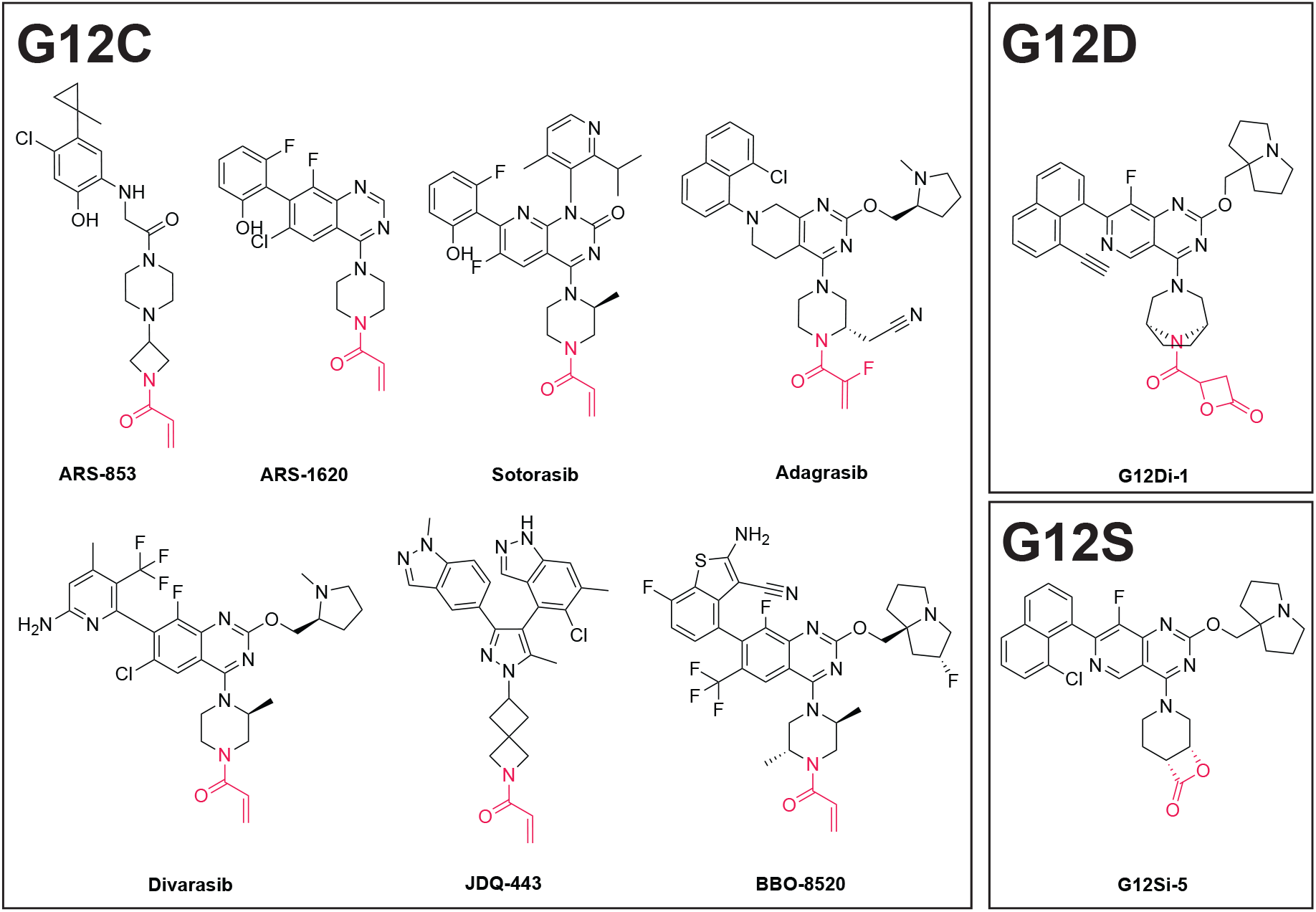
Chemical structures of representative K-Ras(G12C) covalent inhibitors (ARS‐853, ARS‐1620, Sotorasib, Adagrasib, Divarasib, JDQ‐443, and BBO‐8520), K-Ras(G12D) covalent inhibitor (G12Di-1), and K-Ras(G12S) covalent inhibitor (G12Si-5), which were used for structure predictions. Only ARS-853 and ARS-1620 have K-Ras(G12C) co-crystal PDB entries released before the Chai‐1 training cutoff (12 Jan 2021). The warhead is colored red.

**Figure 2.**
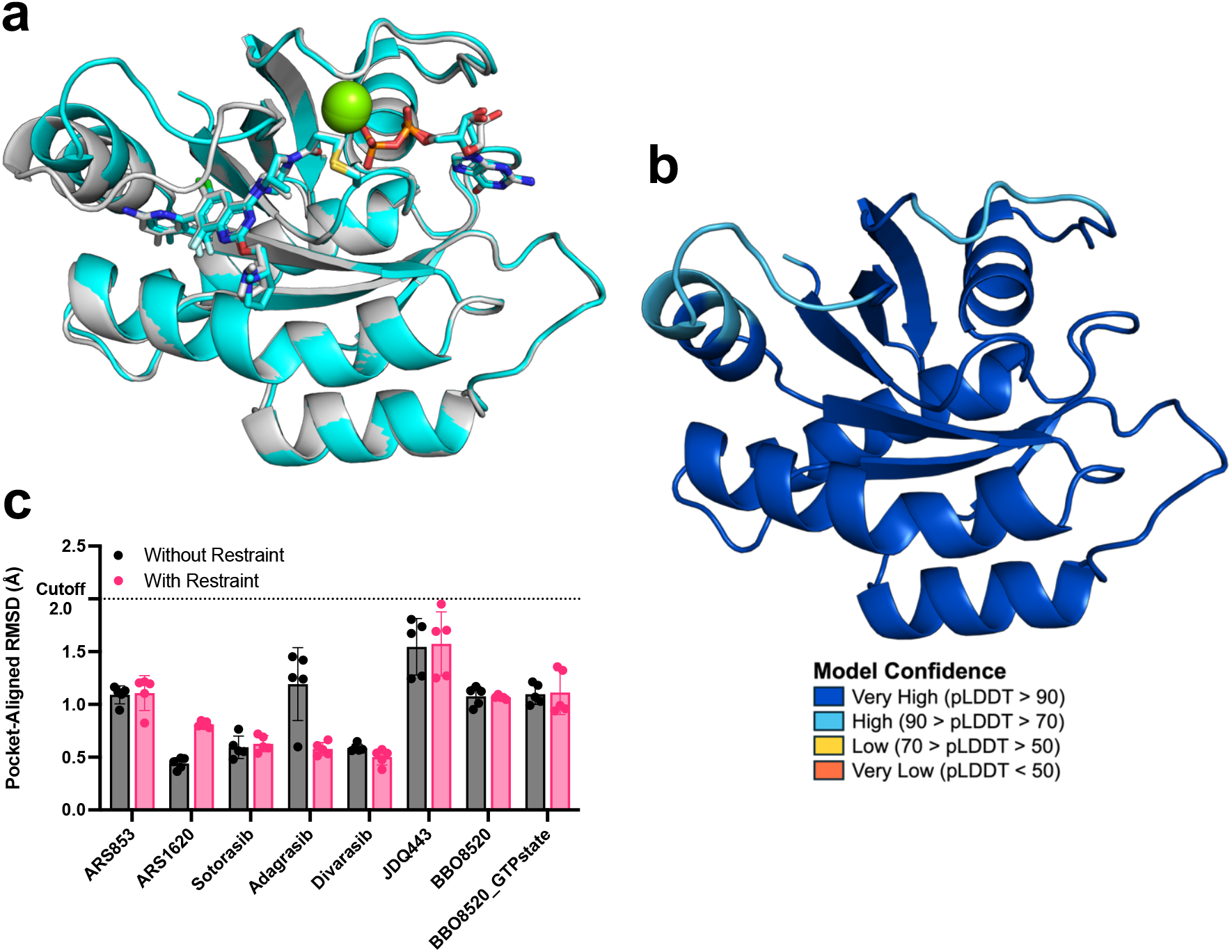
Accurate prediction for covalent K-Ras(G12C) complexes. (a) Overlay of predicted (cyan) vs. crystal (grey) G12C–Divarasib complex; pocket-aligned RMSD 0.56 Å. (b) Predicted K-Ras(G12C)–Divarasib complex colored by per‐residue pLDDT (predicted Local Distance Difference Test); switch-II loop exhibits relatively low confidence score. (c) Pocket-aligned RMSDs for K-Ras(G12C) inhibitors, comparing predictions without and with covalent restraint.

In addition to divarasib, we wondered whether this method could be applied to other covalent K-Ras(G12C) complexes, especially those that were not used as the training set for Chai-1 (training cutoff: 12 Jan 2021). We assembled six additional G12C ligands with diverse chemotypes: ARS-853^21^, ARS-1620^22^, sotorasib, adagrasib, JDQ-443^23^, and BBO-8520^24^ (Figure 1). Chai-1 reproduced all poses within 2.0 Å Pocket-Aligned RMSD, even including JDQ-443, which does not employ the traditional quinazoline scaffold. Remarkably, the model also captured the GTP-bound conformation of BBO-8520, a first-in-class dual-nucleotide-state inhibitor, with 0.99 Å RMSD, comparable to its GDP counterpart (0.95 Å) (Figure 2c). These results demonstrate that single-sequence language-model embeddings are sufficient to accurately position K-Ras(G12C) covalent inhibitors across diverse ligands and nucleotide states, without the need for multiple sequence alignments.

### Covalent Restraint Enables Accurate Modeling of Unconventional K-Ras(G12D) and K-Ras(G12S) Covalent Chemistry

Recent success against K-Ras(G12C) has relied on Michael-acceptor acrylamides that react with the thiolate of Cys12 and dominate existing training sets for structure-prediction and docking algorithms. Pivoting to the more prevalent K-Ras(G12D) mutant^25^ demands electrophiles that can engage an aspartate carboxylate, a markedly less nucleophilic target, which has motivated the discovery of uncommon warheads such as strain-released malolactones^17^, α-diazoacetamide^18^, tunable oxirane^26^, and aziridine^27^. Because only a handful of aspartate-guided covalent complexes have been solved to date, deep-learning models trained primarily on cysteine-reactive ligands lack coverage of these rarer electrophiles.

Benchmark studies show that covalent-docking accuracy falls steeply when the warhead class is under-represented in training data^28,29^, underscoring the possible need to incorporate explicit bond restraints or retraining when tackling Asp-targeting inhibitors. Against this backdrop, we assessed whether Chai-1 could generalize beyond the prevalent cysteine-centric data.

Extending the protocol to the K-Ras(G12D) mutant^25^ failed: in all web-based predictions the β-lactone-based covalent ligand (G12Di-1^17^) was completely displaced from the switch-II pocket (Figure 3a). We suspected that the high precision for G12C predictions might stem from the fact that acrylamide-based, cysteine-directed covalent complexes are substantially overrepresented across the PDB relative to aspartate-directed chemistries, likely biasing the model toward Cys-covalent recognition and geometries thus explaining the discrepancy. However, we did not confine our attention solely to the training data. We considered another factor: the interaction between the nucleophile (amino acid) and the electrophile (warhead). We hypothesized that this displacement could occur because the model typically predicts an energy-minimized local minimum, coupled with the fact that no covalent bond restraint was imposed. As a result, significant steric hindrance between the electrophile and nucleophile prior to covalent bond formation may have led to this displacement. In our G12C case, the relatively small steric clash between acrylamide and cysteine could allow the scaffold’s directing effect to override this, resulting in an accurate prediction, even without a covalent bond restraint. Consistent with this view, a previously reported β-lactone K-Ras complex was crystallized with wild-type K-Ras rather than K-Ras(G12D)^30^, suggesting that the Asp-directed early noncovalent pose is less stable without an explicit covalent restraint.

**Figure 3.**
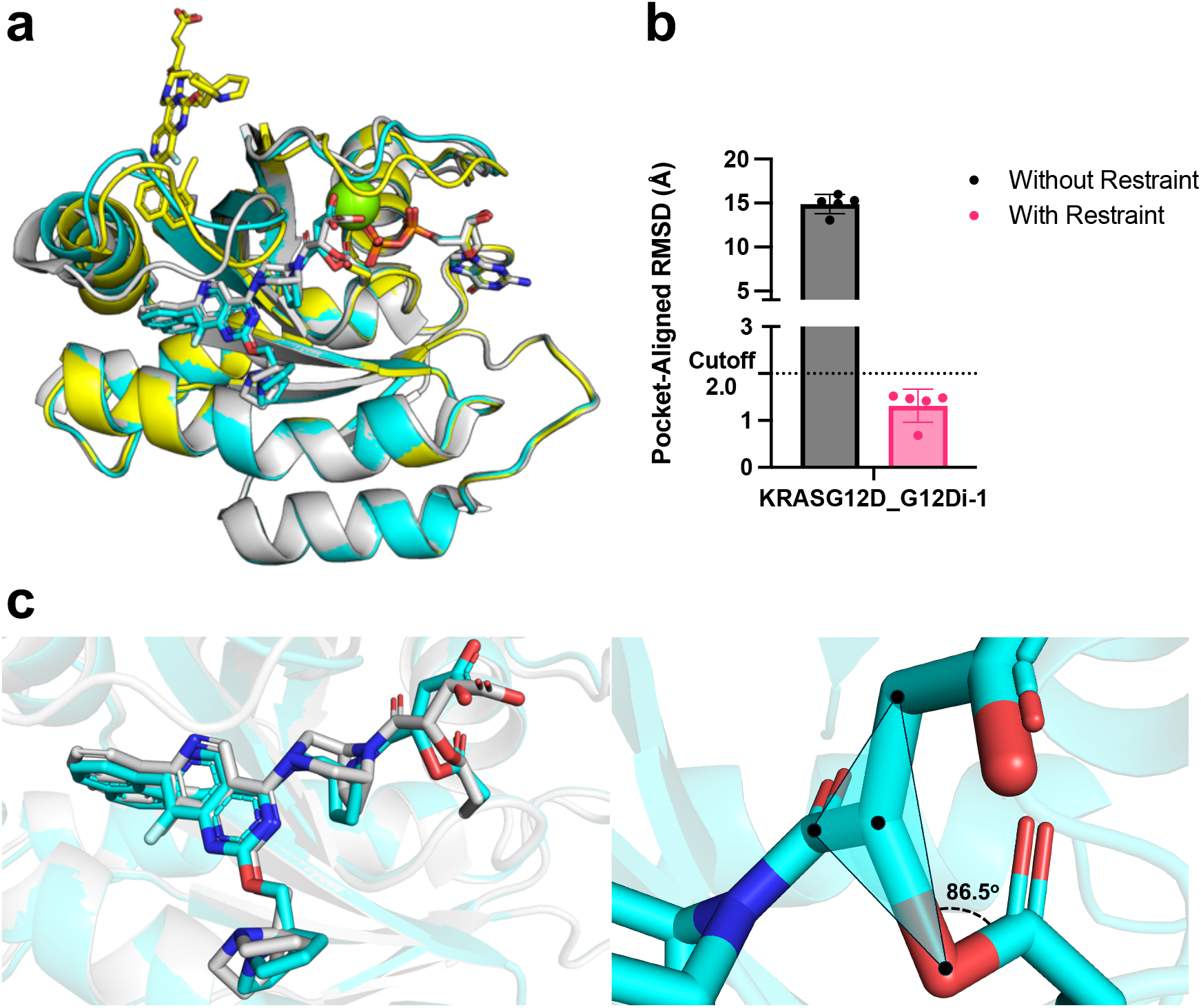
Effect of covalent restraint on K-Ras(G12D) complex. (a) Overlay of naïve single-sequence prediction (yellow) shows ligand ejection from the switch-II pocket. Covalent restraint restores the correct pose (cyan, 0.68 Å Pocket-Aligned RMSD) relative to the crystal structure (grey). (b) Pocket-Aligned RMSD for K-Ras(G12D) inhibitor (G12Di-1), comparing predictions without and with covalent restraint. (c) Close-up overlay of the predicted G12Di-1 structure (cyan) and the crystal structure (grey). The newly formed covalent ester bond exhibits a strained C–O–C angle, and the sp^3^-hybridized carbon within this C–O–C linkage adopts an unusually planar geometry.

To overcome this, we ran the model locally with a covalent bond restraint between G12D and the ligand, which restored the ligand to its experimental pose with a pocket-aligned RMSD of 0.68 Å (Figure 3a, b). However, we obtained a poor consistency across multiple predictions of the switch-II loop region and further, the output is not completely correct in terms of the covalent linker chemistry; the newly formed ester bond has a strained C-O-C angle (measured angle: 86.5°) and an odd planar-like sp^3^-hybridized carbon atom (Figure 3c). For G12C ligands, applying a covalent restraint maintained <2 Å accuracy (Figure 2c).

Encouraged by the success with K‐Ras(G12D), we next tested whether the covalent restraint could similarly enable the accurate prediction of a ligand pose targeting the challenging nucleophile serine in the K‐Ras(G12S) mutant. Serine is also a far less commonly targeted electrophile in covalent drug discovery. For this test, we used G12Si‐5, a β‐lactone–based covalent inhibitor^15^. Without the restraint, the model failed to form the desired covalent bond between the serine hydroxyl and the warhead in all five web-based predictions. However, unlike the complete displacement observed with the G12D inhibitor, the ligand scaffold itself was consistently well-situated within the Switch-II pocket. This suggests the model could predict the binding site accurately but failed to resolve the specific covalent chemistry. Upon applying the covalent restraint, the model successfully recapitulated the experimental binding pose, correctly forming the serine-carbon covalent bond (Figure 4a). Once again, despite this overall accuracy, the predictions revealed a substantial conformational difference within the ligand’s piperidine moiety. Specifically, a near 180° rotation was observed about the C-N bond connecting the piperidine nitrogen to the pyridopyrimidine substituent. Also, the stereochemistry of the piperidine ring was inverted relative to the crystal structure (cis in the experimental pose versus trans in the prediction, Figure 4b). This reflects a known limitation of diffusion-based co-folding models in preserving stereochemical fidelity, highlighting an area for future improvement^31^.

**Figure 4.**
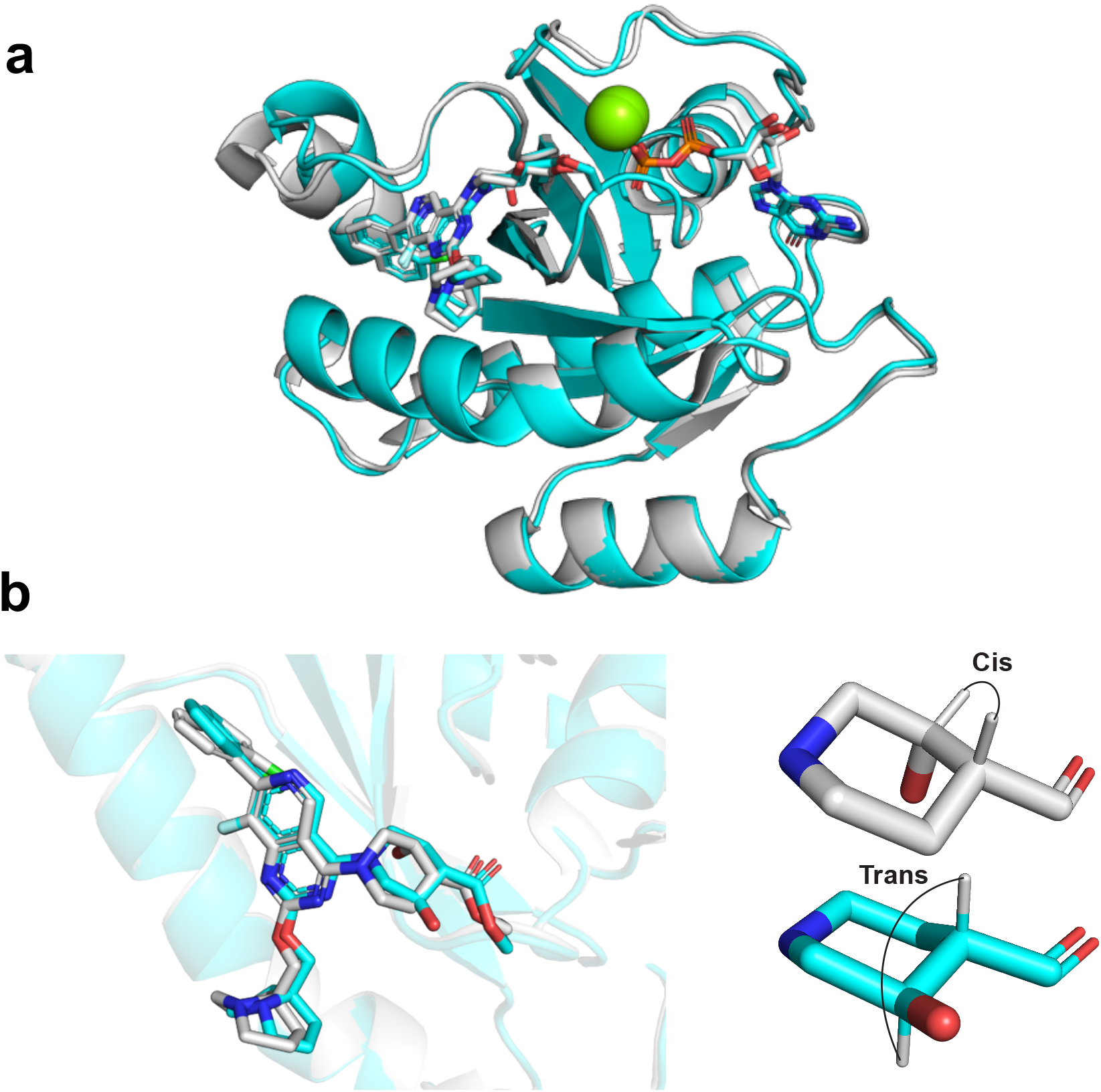
Effect of covalent restraint on K-Ras(G12S) complex. (a) Overlay of the covalently restrained prediction (cyan) and crystallographic pose (grey) of the K-Ras(G12S)– G12Si-5 complex. Without the restraint (not shown), the desired covalent bond is not recapitulated in all five predictions. (b) Close-up overlay of G12Si-5 showing inversion of stereochemistry on the piperidine ring (crystal: cis; prediction: trans).

### Chai-1 Matches AlphaFold3 Pose Accuracy while Delivering ∼40-Fold Higher Throughput in Single-Sequence Mode

To contextualize Chai-1’s computational advantages, we additionally benchmarked AlphaFold3 on the same K-Ras G12 mutant and covalent ligand complexes. We tested AlphaFold3 using MSA and template inputs, widely regarded as state‐of‐the‐art for protein structure prediction due to the rich coevolutionary signal they provide^12^, whereas Chai‐1 was evaluated in single‐sequence mode (i.e., without MSAs). According to the Chai-1 preprint, the model can run in single-sequence mode while preserving most of its performance and markedly reducing runtime^14^. Given this efficiency, Chai-1’s single-sequence mode may be particularly advantageous for large-scale binding pose prediction campaigns involving extensive ligand-protein libraries.

Regarding the ligand–protein pairs predicted earlier by Chai-1, AlphaFold3 achieved a 0.15 ± 0.11 Å lower pocket-aligned RMSD than Chai-1, corresponding to an average improvement of 20 ± 8%. The median paired difference was –0.057 Å. Nevertheless, the difference was not statistically significant (Wilcoxon matched-pairs signed-rank, W = –31, P = 0.074). Importantly, when a 2 Å accuracy cutoff is applied, both models correctly predict all nine ligand poses, yielding 100 % success for this dataset (Figure 5a).

**Figure 5.**
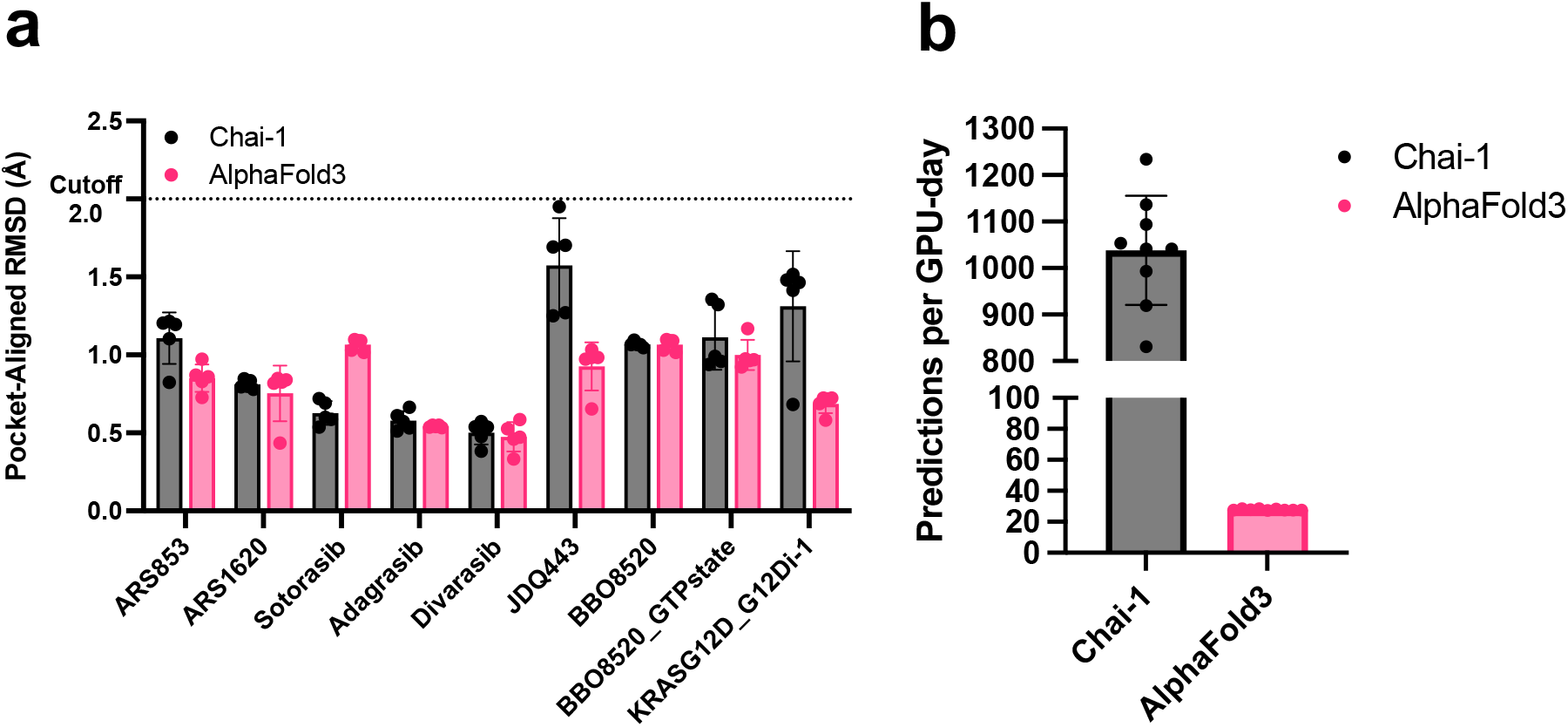
Performance comparison between Chai-1 and AlphaFold3. (a) Pocket-aligned RMSDs for covalent K-Ras(G12C) and K-Ras(G12D) complexes with a covalent restraint. (b) Predicted complexes generated per 24 h on a single A40 GPU for Chai-1 in single-sequence mode versus AlphaFold3 with MSAs and templates.

For computational efficiency, AlphaFold3 required dramatically more time per prediction than Chai-1 in single-sequence mode. Across nine matched K-Ras ligand complexes run on identical A40 (32 GB) GPUs, AF3 consumed ≈ 52 min of CPU wall-time per complex (3,098 ± 48 s), whereas Chai-1 finished in ≈ 1.4 min (84 ± 10 s). Peak RAM usage was modest for both methods (≤ 15 GB), yet AF3’s longer runtime generated a larger memory-seconds integral (1.10 × 10^3^ GB·s vs. 0.87 × 10^3^ GB·s). Extrapolated to a 24-h GPU day, Chai-1 can therefore generate ∼1,000 ligand–protein models, while AF3 is limited to fewer than 30 models (Figure 5b), which is a ∼37-fold reduction in throughput that could pose a substantial challenge for large-scale virtual screening. We acknowledge that most of AF3’s wall-clock time is spent generating MSAs and template features on CPUs rather than on GPU inference; once those files are cached, they can be reused for additional ligands bound to the same protein. However, medicinal-chemistry campaigns rarely focus only on a single invariant sequence. Screening panels of mutants, splice isoforms, or off-targets is routine. Because each new sequence requires a fresh MSA search, AF3’s preprocessing cost re-emerges, whereas Chai-1’s single-sequence mode workflow keeps its short runtime essentially unchanged for typical sequence lengths. In any setting that involves multiple variants or entirely different proteins, Chai-1 retains a decisive, order-of-magnitude efficiency advantage.

## DISCUSSION

Our data validate that Chai-1 can predict covalent inhibitor poses for K-Ras(G12C), K-Ras(G12D), and K-Ras(G12S) at near-crystal resolution, yet several practical constraints remain. First, the model is biased toward the most abundant structures in the PDB and therefore fails to recognize alternative pockets such as the Switch I/II cleft, a transient allosteric groove between Switch-I and Switch-II. In our tests, Chai-1 did not reproduce the published poses for the G12C fragments targeting the Switch I/II cleft (Cpd3) or Switch I (Cpd4)^32^ (Extended Data Fig. 1a, b). These pyridine compounds, which carry an acrylamide warhead, can cross-link Cys12 and occupy the Switch I/II cleft (Cpd3) or Switch I (Cpd4) rather than Switch II; however, Chai-1 predictions placed both Cpd3 and Cpd4 in the Switch II pocket. Another possible explanation could be fragment potency. Cpd3 and Cpd4 are low-affinity fragments (EC50 for MS modification: 23 µM and 11 µM, respectively)^32^, so the model may struggle to stabilize the alternative pocket without a tight binding, fully optimized ligand. Potent, chemically optimized ligands impose stronger geometric constraints, guiding the model toward the correct binding mode. Thus, it would be worthwhile to test Chai-1 with higher-affinity Switch I/II ligands^33,34^ to determine the exact cause.

Second, the model struggles with fine-grained chemical details. For the K-Ras(G12D) complex, the predicted covalent adduct displays improper bond angles (Figure 3c). A possible corrective workflow would be to generate an initial prediction and then run a short MD simulation or a Rosetta FastRelax^35^ to obtain a more physically relevant structure. For the G12S mutant and its inhibitor G12Si-5, Chai-1 with a covalent bond restraint largely reproduced the binding pocket and ligand orientation (Figure 4a). However, it did not preserve the correct stereochemistry (Figure 4b) and missed a key hydrogen bond network in the ground truth structure^15^, even though the input ligand was supplied with the proper stereochemical specification. This shortcoming likely stems from the network’s inability to lock stereochemistry for reliable conformational analysis or to model reaction mechanisms and explicit crystallographic water molecules, which often stabilize the true, lower-energy state through hydrogen bonding. We regard this as a fundamental limitation of deep-learning structure predictors such as Chai-1 and AF3. Encouragingly, newer models have been developed to address these gaps. For example, Boltz-2x^36,37^ adds steering potentials during inference and joint learning for structure and affinity to push poses toward physically consistent geometry. Separately, DiffDock-Glide^38^ refines generative samples with Glide minimization to enforce local stereochemistry and relieve clashes. Finally, MIC^39^ assigns positions for water and ions using deep learning to help restore stabilizing hydration networks.

Third, no current deep learning-based structure prediction tool automatically removes the leaving group after a covalent bond is formed. Our workaround for G12C inhibitors is to model acrylamide as propanamide; for G12Di-1 and G12Si-5, we input the ligand SMILES in its “post-reaction” form. Ideally, the software would apply covalent constraints that mimic authentic adducts, but that capability is not yet available. The challenge is even greater for arginine-specific warheads, which form two covalent bonds with a single residue^16^; to our knowledge, no existing pipeline can handle that chemistry. As an alternative, treating the ligand-modified residue as a non-canonical amino acid and predicting the covalently modified protein without an explicit ligand entry is possible, but the approach has not been optimized for high-throughput covalent docking.

Fourth, our validation set is limited to K-Ras G12 point mutants. Broader generality across other targets, including additional K-Ras cysteine variants (G13C, G61C) or even H- and N-Ras, remains untested. Finally, while Chai-1 is faster than MSA-based methods, each prediction still requires ∼1–2 min of GPU time, making large-library screening slower and costlier than conventional docking. Even so, Chai-1’s fully open, scriptable architecture should enable the community to iteratively close these gaps, positioning it as a routine and scalable engine for covalent lead optimization.

## METHODS

### Chai-1 web predictions

All jobs were submitted to the Chai-1 web portal at http://lab.chaidiscovery.com/. Predictions ran in single-sequence mode without using multiple sequence alignments, template structures, or any restraints.

### Local GPU predictions with covalent restraint

GPU runs were carried out on the UCSF Wynton HPC (qb3-atgpu queue) using NVIDIA A40 GPUs. The source codes were drawn from the following GitHub repositories (Chai-1: https://github.com/chaidiscovery/chai-lab, AlphaFold3: https://github.com/google-deepmind/alphafold3). For ligand preparation, we used the SMILES string of the ligand as present in the crystal structure (after the covalent bond-forming reaction), with the leaving group already removed.

### RMSD calculation

Pocket-aligned RMSDs were computed following the procedure described in the AlphaFold3 Supporting Information^12^. Briefly, the binding pocket is defined as all heavy-atom coordinates located within 10 Å of any heavy atom of the ligand in the reference crystal structure. The predicted model is then superposed onto the ground-truth structure by least-squares rigid-body alignment using only the Cα atoms of residues within this pocket. Finally, the RMSD is calculated over all heavy atoms of the ligand. A reproducible PyMOL/Python workflow and scripts are provided (alignment to a pocket selection, ligand extraction from ground truth and predictions, and RDKit-based RMSD with optional manual atom-pair overrides), together with example commands. Code and exact commands are included in the SI.

## Supporting information

Supplementary_Data.zip: inputs/outputs, PyMOL sessions, and pocket-aligned RMSD workflow

## SUPPORTING INFORMATION

A single compressed archive, **Supplementary_Data.zip**, accompanies this Article. It contains three top-level directories:

### 1_Without Covalent Restraint_Web/

Chai-1 single-sequence web runs (no covalent restraint).

- **Chai-1 Inputs/** – protein sequence and ligand SMILES (FASTA files) used for inference.
- **Predicted Structures/** – raw model outputs (e.g., pred.model_idx_0 … pred.model_idx_4, in CIF as produced by the service).
- **PyMOL Sessions/** –.pse files showing the alignment of five predictions (single seed) to the reference structure for visualization.

### 2_With Covalent Restraint/

Local runs with explicit covalent restraints.

- **AlphaFold3/** – input files (JSON + ligand CIF), the corresponding predicted structures, and.pse files showing the alignment.
- **Chai-1/** – input files (FASTA + restraint file), the corresponding predicted structures, and.pse files showing the alignment.

### 3_Pocket-Aligned_RMSD/

Reproducible workflow, scripts, and raw results for the pocket-aligned RMSD procedure.

- **README.md** – step-by-step instructions (pocket definition in PyMOL, alignment, ligand extraction, and RMSD computation) consistent with the Methods text.
- **predictions_align_to_pocket.py** *(used in PyMOL)* – aligns pred.model_idx_0 … pred.model_idx_4 to the ground-truth **pocket** using only pocket **Cα** atoms.
- **save_ligands.py** *(used in PyMOL)* – extracts the ground-truth ligand (gt_ligand.pdb) and predicted ligands (pred.model_idx_#_ligand.pdb) based on user-set selections (edit gt_selection/selection inside the script).
- **pocket_aligned_ligand_RMSD.py** *(Python; RDKit + NumPy)* – computes heavy-atom ligand RMSDs between the reference and each pocket-aligned prediction; supports optional **manual atom-pair overrides**; writes a CSV summary with sanity checks.
- **Raw Pocket-Aligned RMSDs and Prediction Throughputs.xlsx** – raw RMSD values for all predictions and the measured prediction throughput figures used in the paper.

## Author Contributions

CRediT: **Sungwon Jung**: Conceptualization, Data curation, Formal analysis, Investigation, Methodology, Validation, Visualization, Writing – original draft, Writing – review & editing; **Qinheng Zheng**: Investigation (suggested experiments), Writing – review & editing; **Kevan M. Shokat**: Conceptualization, Funding acquisition, Resources, Supervision, Writing – review & editing.

## ACKNOWLEDGMENTS

Portions of this work were performed on the Wynton HPC Co-Op cluster, which is supported by UCSF research faculty and UCSF institutional funds. The authors wish to thank the UCSF Wynton team for their ongoing technical support of the Wynton environment. K.M.S. acknowledges support from the NIH (5R01CA244550), a gift from the Sjöberg Foundation, and the LRRK2 Investigative Therapeutics Exchange (LITE) program of the Michael J. Fox Foundation. S.J. acknowledges support from the Fulbright U.S.–Korea Presidential STEM Initiative Award, administered by the Korean American Educational Commission (Fulbright Korea). Q.Z. is the Connie and Bob Lurie Fellow of the Damon Runyon Cancer Research Foundation (DRG-2434-21) and acknowledges support from the UCSF Pancreas Center via a Mentored Scientist Award.

## EXTENDED DATA FIGURES

**Extended Data Figure 1.**
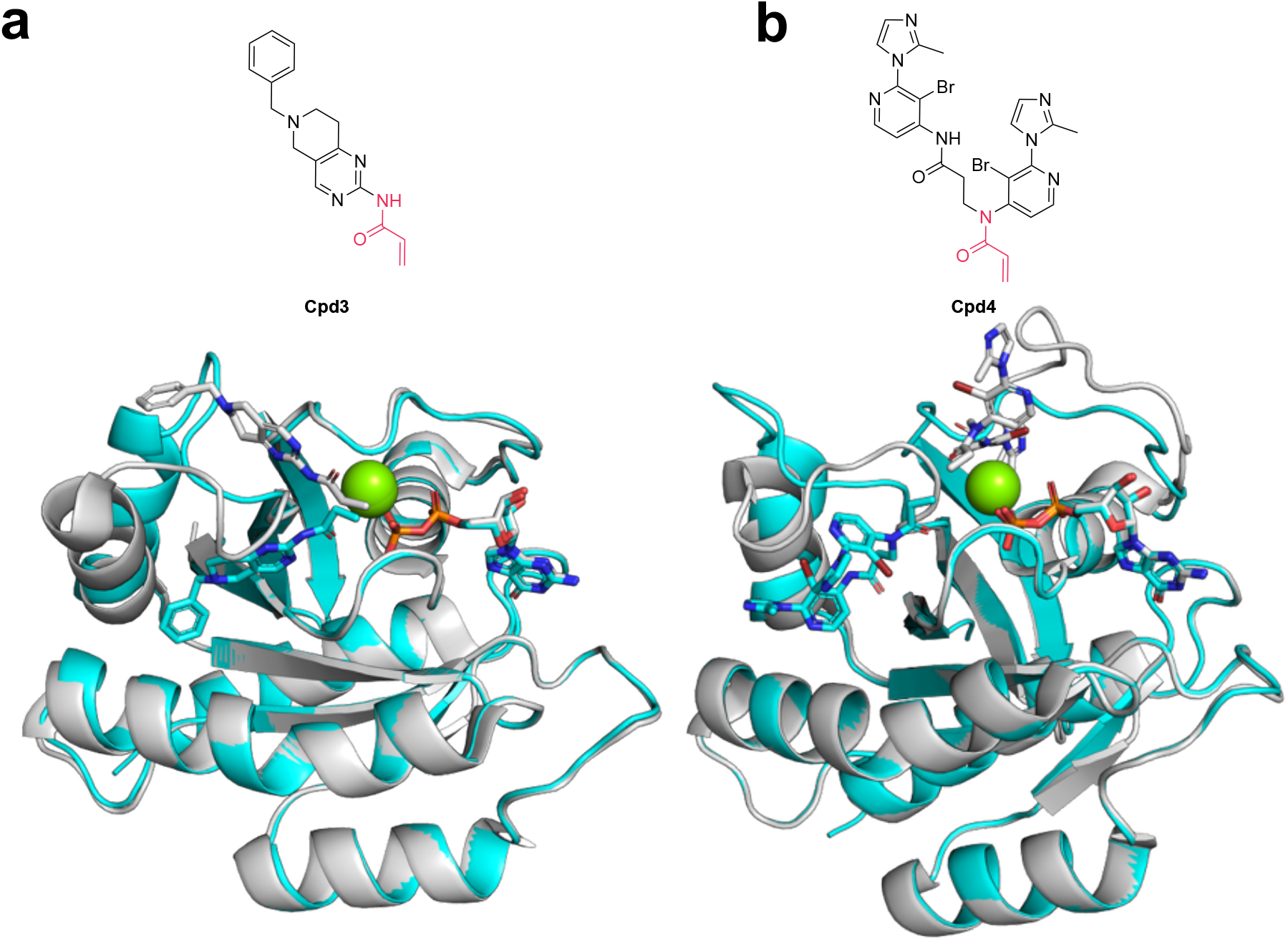
Lack of pose reproducibility for Switch I and Switch I/II-pocket fragments. (a) Chemical structure of Cpd3 and overlay of the predicted (cyan) versus crystallographic (grey) KRAS (G12C)–Cpd3 complex. (b) Chemical structure of Cpd4 and overlay of the predicted versus crystallographic poses. In both cases, the model places the ligands in the Switch II pocket rather than the intended Switch I/II cleft (Cpd3) or Switch I (Cpd4), respectively.

