## Supplementary_Data.zip: inputs/outputs, PyMOL sessions, and pocket-aligned RMSD workflow for "Single-sequence deep learning delivers crystal-quality models of covalent K-Ras G12 hotspot complexes": TERMS_OF_USE.md

# ALPHAFOLD 3 OUTPUT TERMS OF USE

Last Modified: 2024-11-09

By using AlphaFold 3 Output (as defined below), without having agreed to
[AlphaFold 3 Model Parameters Terms of Use](https://github.com/google-deepmind/alphafold3/blob/main/WEIGHTS_TERMS_OF_USE.md),
you agree to be bound by these AlphaFold 3 Output Terms of Use between you (or
your organization, as applicable) and Google LLC (these "**Terms**").

If you are using Output on behalf of an organization, you confirm you are
authorized either explicitly or implicitly to agree to, and are agreeing to,
these Terms as an employee on behalf of, or otherwise on behalf of, your
organization.

If you have agreed to
[AlphaFold 3 Model Parameters Terms of Use](https://github.com/google-deepmind/alphafold3/blob/main/WEIGHTS_TERMS_OF_USE.md),
your use of Output are governed by those terms. **If you have not agreed to
[AlphaFold 3 Model Parameters Terms of Use](https://github.com/google-deepmind/alphafold3/blob/main/WEIGHTS_TERMS_OF_USE.md)
and do not agree to these Terms, do not use Output or permit any third party to
do so on your behalf.**

When we say "**you**", we mean the individual or organization using Output. When
we say "**we**", "**us**" or "**Google**", we mean the entities that belong to
the Google group of companies, which means Google LLC and its affiliates.

## Key Definitions

As used in these Terms:

"**AlphaFold 3**" means the AlphaFold 3 Code and Model Parameters.

"**AlphaFold 3 Code**" means the AlphaFold 3 source code: (a) identified at
[public GitHub repo](https://github.com/google-deepmind/alphafold3/), or such
other location in which we may make it available from time to time, regardless
of the source that it was obtained from; and (b) made available by Google to
organizations for their use in accordance with the
[AlphaFold 3 Model Parameters Terms of Use](https://github.com/google-deepmind/alphafold3/blob/main/WEIGHTS_TERMS_OF_USE.md)
(not these Terms) together with (i) modifications to that code, (ii) works based
on that code, or (iii) other code or machine learning model which incorporates,
in full or in part, that code.

"**Model Parameters**" means the trained model weights and parameters made
available by Google to organizations (at its sole discretion) for their use in
accordance with the
[AlphaFold 3 Model Parameters Terms of Use](https://github.com/google-deepmind/alphafold3/blob/main/WEIGHTS_TERMS_OF_USE.md)
(not these Terms), together with (a) modifications to those weights and
parameters, (b) works based on those weights and parameters, or (c) other code
or machine learning model which incorporates, in full or in part, those weights
and parameters.

"**Output**" means the structure predictions and all related information
provided by AlphaFold 3, together with any visual representations, computational
predictions, descriptions, modifications, copies, or adaptations that are
substantially derived from Output.

## Use restrictions

[AlphaFold 3](https://blog.google/technology/ai/google-deepmind-isomorphic-alphafold-3-ai-model/)
belongs to us. Output are made available free of charge, for non-commercial use
only, in accordance with the following use restrictions. You must not use nor
allow others to use Output:

1.  **On behalf of a commercial organization or in connection with any
    commercial activities, including research on behalf of commercial
    organizations.**

    1.  This means that only non-commercial organizations (*i.e.*, universities,
        non-profit organizations and research institutes, educational,
        journalism and government bodies) may use Output for their
        non-commercial activities. Output are not available for use by any other
        types of organization, even if conducting non-commercial work.

    2.  If you are a researcher affiliated with a non-commercial organization,
        provided **you are not a commercial organisation or acting on behalf of
        a commercial organisation**, you can use Output for your non-commercial
        affiliated research.

    3.  You must not share Output with any commercial organization. The only
        exception is making Output publicly available (including, indirectly, to
        commercial organizations) via a scientific publication or open source
        release or using these to support journalism, each of which are
        permitted.

2.  **To misinform, misrepresent or mislead**, including:

    1.  providing false or inaccurate information in relation to your access to
        or use of Output;

    2.  misrepresenting your relationship with Google - including by using
        Google’s trademarks, trade names, logos or suggesting endorsement by
        Google without Google’s permission to do so - nothing in these Terms
        grants such permission;

    3.  misrepresenting the origin of Output;

    4.  distributing misleading claims of expertise or capability, or engaging
        in the unauthorized or unlicensed practice of any profession,
        particularly in sensitive areas (*e.g.*, health); or

    5.  making decisions in domains that affect material or individual rights or
        well-being (*e.g.*, healthcare).

3.  **To perform, promote or facilitate dangerous, illegal or malicious
    activities**, including:

    1.  promoting or facilitating the sale of, or providing instructions for
        synthesizing or accessing, illegal substances, goods or services;

    2.  abusing, harming, interfering, or disrupting any services, including
        generating or distributing content for deceptive or fraudulent
        activities or malware;

    3.  generating or distributing any content that infringes, misappropriates,
        or otherwise violates any individual’s or entity’s rights (including,
        but not limited to rights in copyrighted content); or

    4.  attempting to circumvent these Terms.

4.  **To train or create machine learning models or related technology for
    biomolecular structure prediction similar to AlphaFold 3 as made available
    by Google ("Derived Models"),** including via distillation or other
    methods**.** For the avoidance of doubt, the use restrictions set out in
    these Terms would apply in full to any Derived Models created in breach of
    these Terms.

5.  **Without providing conspicuous notice that published or distributed Output
    is provided under and subject to these Terms and of any modifications you
    make to Output.**

    1.  This means if you remove, or cause to be removed (for example by using
        third-party software), these Terms, or any notice of these Terms, from
        Output, you must ensure further distribution or publication is
        accompanied by a copy of the
        [AlphaFold 3 Output Terms of Use](https://github.com/google-deepmind/alphafold3/blob/main/OUTPUT_TERMS_OF_USE.md)
        and a "*Legally Binding Terms of Use*" text file that contains the
        following notice:

        "*By using this information, you agree to AlphaFold 3 Output Terms of
        Use found at
        https://github.com/google-deepmind/alphafold3/blob/main/OUTPUT_TERMS_OF_USE.md.*

        *To request access to the AlphaFold 3 model parameters, follow the
        process set out at https://github.com/google-deepmind/alphafold3. You
        may only use these if received directly from Google. Use is subject to
        terms of use available at
        https://github.com/google-deepmind/alphafold3/blob/main/WEIGHTS_TERMS_OF_USE.md.*"

    2.  You must not include any additional or different terms that conflict
        with the
        [AlphaFold 3 Output Terms of Use](https://github.com/google-deepmind/alphafold3/blob/main/OUTPUT_TERMS_OF_USE.md).

6.  **Distribute Output, or disclose findings arising from using AlphaFold 3
    without citing our paper:** [Abramson, J et al. Accurate structure
    prediction of biomolecular interactions with AlphaFold 3. *Nature*
    (2024)](https://www.nature.com/articles/s41586-024-07487-w). For the
    avoidance of doubt, this is an additional requirement to the notice
    requirements set out above.

We grant you a non-exclusive, royalty-free, revocable, non-transferable and
non-sublicensable (except as expressly permitted in these Terms) license to any
intellectual property rights we have in Output to the extent necessary for these
purposes. You agree that your right to use and share Output is subject to your
compliance with these Terms. If you breach these Terms, Google reserves the
right to request that you delete and cease use or sharing of Output in your
possession or control and prohibit you from using the AlphaFold 3 Assets
(including as made available via
[AlphaFold Server](https://alphafoldserver.com/about)). You agree to immediately
comply with any such request.

## Disclaimers

Nothing in these Terms restricts any rights that cannot be restricted under
applicable law or limits Google’s responsibilities except as allowed by
applicable law.

**Output are provided on an "as is" basis, without warranties or conditions of
any kind, either express or implied, including any warranties or conditions of
title, non-infringement, merchantability, or fitness for a particular purpose.
You are solely responsible for determining the appropriateness of using or
distributing any of the Output and assume any and all risks associated with your
use or distribution of any Output and your exercise of rights and obligations
under these Terms. You and anyone you share Output with are solely responsible
for these and their subsequent uses.**

**Output are predictions with varying levels of confidence and should be
interpreted carefully. Use discretion before relying on, publishing, downloading
or otherwise using Output.**

**Output are for theoretical modeling only. These are not intended, validated,
or approved for clinical use. You should not use these for clinical purposes or
rely on them for medical or other professional advice. Any content regarding
those topics is provided for informational purposes only and is not a substitute
for advice from a qualified professional.**

## Liabilities

To the extent allowed by applicable law, you will indemnify Google and its
directors, officers, employees, and contractors for any third-party legal
proceedings (including actions by government authorities) arising out of or
relating to your unlawful use of Output or violation of these Terms. This
indemnity covers any liability or expense arising from claims, losses, damages,
judgments, fines, litigation costs, and legal fees, except to the extent a
liability or expense is caused by Google's breach, negligence, or willful
misconduct. If you are legally exempt from certain responsibilities, including
indemnification, then those responsibilities don’t apply to you under these
terms.

In no circumstances will Google be responsible for any indirect, special,
incidental, exemplary, consequential, or punitive damages, or lost profits of
any kind, even if Google has been advised of the possibility of such damages.
Google’s total, aggregate liability for all claims arising out of or in
connection with these Terms or Output, including for its own negligence, is
limited to $500.

## Governing law and disputes

These Terms will be governed by the laws of the State of California. The state
or federal courts of Santa Clara County, California shall have exclusive
jurisdiction of any dispute arising out of these Terms.

Given the nature of scientific research, it may take some time for any breach of
these Terms to become apparent. To the extent allowed by applicable law, any
legal claims relating to these Terms or Output can be initiated until the later
of (a) the cut-off date under applicable law for bringing the legal claim; or
(b) two years from the date you or Google (as applicable) became aware, or
should reasonably have become aware, of the facts giving rise to that claim. You
will not argue limitation, time bar, delay, waiver or the like in an attempt to
bar an action filed within that time period, and neither will we.

All rights not specifically and expressly granted to you by these Terms are
reserved to Google. No delay, act or omission by Google in exercising any right
or remedy will be deemed a waiver of any breach of these Terms and Google
expressly reserves any and all rights and remedies available under these Terms
or at law or in equity or otherwise, including the remedy of injunctive relief
against any threatened or actual breach of these Terms without the necessity of
proving actual damages.

## Miscellaneous

Google may update these Terms (1) to reflect changes in how it does business,
(2) for legal, regulatory or security reasons, or (3) to prevent abuse or harm.
The version of these Terms that were effective on the date the relevant Output
was generated will apply to your use of that Output.

If it turns out that a particular provision of these Terms is not valid or
enforceable, this will not affect any other provisions.
