## Supplementary_Data.zip: inputs/outputs, PyMOL sessions, and pocket-aligned RMSD workflow for "Single-sequence deep learning delivers crystal-quality models of covalent K-Ras G12 hotspot complexes": pocket-aligned_ligand_RMSD.py

```
#!/usr/bin/env python

# -----------------------------------------------------------------------------
# Sungwon Jung
# -----------------------------------------------------------------------------

# This script is a conversion of a Jupyter Notebook for calculating RMSD between a ground truth ligand and multiple predicted ligand poses, with optional manual overrides for atom pairings.
# It uses RDKit for molecular handling and NumPy for numerical calculations.
# The script processes PDB files, computes RMSD values, performs sanity checks, and outputs results in CSV format.
# Manual overrides can be applied to adjust atom pairings for more accurate RMSD calculations.

# --- Dependencies ---
# Before running, make sure you have the required Python libraries installed.
# You can install them using pip:
# pip install rdkit-pypi
# pip install "numpy<2"
# Run with overrides: `python multi_pocket_ligand_RMSD_AF3.py --use-overrides` (omit the flag to disable)

import os
import re
import csv
import numpy as np
from rdkit import Chem
from rdkit.Chem import AllChem
import argparse

# --- CLI options ---
parser = argparse.ArgumentParser(description="Compute pocket-aligned RMSDs with optional manual overrides.")
parser.add_argument("--use-overrides", action="store_true", help="Apply manual atom pairing overrides defined in the script.")
args = parser.parse_args()
APPLY_OVERRIDES = args.use_overrides

# ==============================================================================
# Step 1: Prepare the Template Molecule from SMILES
# ==============================================================================
print("--- Step 1: Preparing Template Molecule ---")
# Provided SMILES (original connectivity before conversion to single bonds)
smiles_original = "Fc1c(Oc2ccc3c(B(O)OC3)c2)ccc(CN)c1"

# Convert SMILES to an RDKit molecule
mol_template = Chem.MolFromSmiles(smiles_original)
if mol_template is None:
    raise ValueError("Error converting SMILES to molecule.")
# Add explicit hydrogens
mol_template = Chem.AddHs(mol_template)

# Convert all bonds to SINGLE bonds to remove connectivity complications
for bond in mol_template.GetBonds():
    bond.SetBondType(Chem.rdchem.BondType.SINGLE)

# Generate and print the canonical SMILES
canonical_smiles = Chem.MolToSmiles(mol_template, canonical=True)
print(f"Canonical SMILES (all single bonds): {canonical_smiles}\n")


# ==============================================================================
# Step 2: Load Ground Truth and Predicted PDB Files
# ==============================================================================
print("--- Step 2: Loading PDB Files ---")
# Load the ground truth molecule
gt_pdb_file = "gt_ligand.pdb"
if not os.path.exists(gt_pdb_file):
    raise FileNotFoundError(f"Ground truth file not found: {gt_pdb_file}")
gt_mol = Chem.MolFromPDBFile(gt_pdb_file, removeHs=False)
gt_mol = Chem.AddHs(gt_mol)
print(f"Loaded ground truth: {gt_pdb_file}")


# Load multiple prediction molecules (up to 5)
pred_mols = []
pred_file_patterns = [f"pred.model_idx_{i}_ligand.pdb" for i in range(5)]

for i, pred_file in enumerate(pred_file_patterns):
    if os.path.exists(pred_file):
        pred_mol = Chem.MolFromPDBFile(pred_file, removeHs=False)
        pred_mol = Chem.AddHs(pred_mol)
        pred_mols.append((i, pred_mol))
        print(f"Loaded prediction {i}: {pred_file}")
    else:
        print(f"Warning: Could not find prediction file {pred_file}")

if not pred_mols:
    raise ValueError("No prediction files found! Please check file paths.")
print("")


# ==============================================================================
# Step 3: Find Atom Mappings using a Substructure Match
# ==============================================================================
print("--- Step 3: Finding Atom Mappings ---")
# Use the template molecule (with single bonds) to find corresponding atom indices
mapping_gt = gt_mol.GetSubstructMatch(mol_template)
if not mapping_gt:
    raise ValueError("Substructure matching failed for ground truth molecule.")

# Get mappings for all prediction molecules
pred_mappings = []
for idx, pred_mol in pred_mols:
    mapping_pred = pred_mol.GetSubstructMatch(mol_template)
    if not mapping_pred:
        print(f"Warning: Substructure matching failed for prediction {idx}. Skipping.")
        continue
    pred_mappings.append((idx, pred_mol, mapping_pred))

print(f"Found {len(pred_mappings)} valid prediction molecules with mappings.")
print(f"Mapping for ground truth molecule has {len(mapping_gt)} atoms\n")


# ==============================================================================
# Step 4: Define Helper Functions for RMSD Calculation and Reporting
# ==============================================================================

def compute_and_report(gt_mol, mapping_gt, pred_idx, pred_mol, mapping_pred, manual_overrides=None, suffix=""):
    """
    Computes coordinates, RMSD, sanity checks, and outputs CSV files.
    """
    # Apply manual overrides if provided
    if manual_overrides:
        # Create a new copy of the predicted mapping as a list to apply overrides
        mapping_pred_override = list(mapping_pred)
        for csv_idx, new_pred_orig_idx in manual_overrides.items():
            if csv_idx <= len(mapping_pred_override):
                old_val = mapping_pred_override[csv_idx - 1]
                mapping_pred_override[csv_idx - 1] = new_pred_orig_idx - 1  # Convert from 1-based to 0-based
                print(f"Overriding pairing for ordering index {csv_idx}: "
                      f"Predicted atom changed from {old_val+1} to {new_pred_orig_idx}")
            else:
                print(f"Warning: ordering index {csv_idx} is out of range for the predicted mapping.")
        mapping_pred = mapping_pred_override

    # Get conformers
    conf_gt = gt_mol.GetConformer()
    conf_pred = pred_mol.GetConformer()

    # Extract coordinates in the order given by the mapping
    coords_gt = np.array([list(conf_gt.GetAtomPosition(idx)) for idx in mapping_gt])
    coords_pred = np.array([list(conf_pred.GetAtomPosition(idx)) for idx in mapping_pred])

    # Output coordinates to CSV files
    gt_filename = f"gt_coordinates{suffix}.csv"
    pred_filename = f"pred{pred_idx}_coordinates{suffix}.csv"
    with open(gt_filename, "w", newline='') as f:
        writer = csv.writer(f)
        writer.writerow(["atom_idx", "orig_idx", "atom_type", "x", "y", "z"])
        for i, idx in enumerate(mapping_gt):
            atom = gt_mol.GetAtomWithIdx(idx)
            pos = conf_gt.GetAtomPosition(idx)
            writer.writerow([i + 1, idx + 1, atom.GetSymbol(), f"{pos.x:.4f}", f"{pos.y:.4f}", f"{pos.z:.4f}"])
    with open(pred_filename, "w", newline='') as f:
        writer = csv.writer(f)
        writer.writerow(["atom_idx", "orig_idx", "atom_type", "x", "y", "z"])
        for i, idx in enumerate(mapping_pred):
            atom = pred_mol.GetAtomWithIdx(idx)
            pos = conf_pred.GetAtomPosition(idx)
            writer.writerow([i + 1, idx + 1, atom.GetSymbol(), f"{pos.x:.4f}", f"{pos.y:.4f}", f"{pos.z:.4f}"])
    print(f"Coordinate files saved as '{gt_filename}' and '{pred_filename}'")

    # Compute RMSD
    rmsd = np.sqrt(np.mean(np.sum((coords_gt - coords_pred)**2, axis=1)))
    print(f"Prediction {pred_idx} - Computed RMSD: {rmsd:.4f}")

    # Sanity Checks
    if len(coords_gt) != len(coords_pred):
        print(f"WARNING: Different number of atoms! GT: {len(coords_gt)}, Pred: {len(coords_pred)}")
    else:
        print(f"Sanity check passed: Both coordinate sets have {len(coords_gt)} atoms")

    matching_atoms = True
    for i, (gt_idx, pred_idx) in enumerate(zip(mapping_gt, mapping_pred)):
        gt_atom = gt_mol.GetAtomWithIdx(gt_idx)
        pred_atom = pred_mol.GetAtomWithIdx(pred_idx)
        if gt_atom.GetSymbol() != pred_atom.GetSymbol():
            matching_atoms = False
            print(f"WARNING: Atom type mismatch at position {i+1}: {gt_atom.GetSymbol()} vs {pred_atom.GetSymbol()}")
    if matching_atoms:
        print("Sanity check passed: All atom types match in the same order")

    # Per-atom distance analysis
    atom_distances = np.sqrt(np.sum((coords_gt - coords_pred)**2, axis=1))
    print(f"Average atomic distance: {np.mean(atom_distances):.4f} Å")
    print(f"Maximum atomic distance: {np.max(atom_distances):.4f} Å at atom index {np.argmax(atom_distances)+1}")

    # Top 10 atoms with largest differences
    print("\nAtoms with largest position differences:")
    sorted_indices = np.argsort(atom_distances)[::-1]
    for rank in range(min(10, len(sorted_indices))):
        idx = sorted_indices[rank]
        gt_orig_idx = mapping_gt[idx] + 1
        pred_orig_idx = mapping_pred[idx] + 1
        gt_atom = gt_mol.GetAtomWithIdx(mapping_gt[idx])
        print(f"  {rank+1}. GT atom_idx: {idx+1} (orig_idx: {gt_orig_idx}), "
              f"Pred atom_idx: {idx+1} (orig_idx: {pred_orig_idx})")
        print(f"     Atom {idx+1} ({gt_atom.GetSymbol()}): Distance = {atom_distances[idx]:.4f} Å")
        print(f"     GT   position: ({coords_gt[idx][0]:.4f}, {coords_gt[idx][1]:.4f}, {coords_gt[idx][2]:.4f})")
        print(f"     Pred position: ({coords_pred[idx][0]:.4f}, {coords_pred[idx][1]:.4f}, {coords_pred[idx][2]:.4f})")
    print("\n")

    return rmsd

def process_prediction_with_overrides(gt_mol, mapping_gt, pred_idx, pred_mol, mapping_pred, manual_overrides_dict=None):
    """
    Process a single prediction with possible manual overrides.
    """
    print(f"\n### Processing Prediction {pred_idx} ###")

    # First run without overrides
    print(f"--- First Run for Prediction {pred_idx}: Original Pairings ---")
    base_rmsd = compute_and_report(gt_mol, mapping_gt, pred_idx, pred_mol, mapping_pred,
                                   suffix=f"_pred{pred_idx}")

    # Second run with overrides if they exist
    if manual_overrides_dict and pred_idx in manual_overrides_dict:
        print(f"\n--- Second Run for Prediction {pred_idx}: With Manual Overrides ---")
        override_rmsd = compute_and_report(gt_mol, mapping_gt, pred_idx, pred_mol, mapping_pred,
                                          manual_overrides=manual_overrides_dict[pred_idx],
                                          suffix=f"_pred{pred_idx}_override")
        return base_rmsd, override_rmsd
    else:
        return base_rmsd, base_rmsd # Return base_rmsd for both if no override

# ==============================================================================
# Step 5: Main Execution - Process All Predictions
# ==============================================================================
print("--- Step 5: Processing All Predictions and Calculating RMSD ---")
print(f"Manual overrides are {'ENABLED' if APPLY_OVERRIDES else 'DISABLED'}.")

# Dictionary for manual overrides
# Format: {pred_idx: {ordering_idx: new_pred_orig_idx, ...}}
manual_overrides_dict = {
    0: {31: 38, 32: 37, 33: 36, 35: 35, 36: 34, 37: 33},
    1: {31: 38, 32: 37, 33: 36, 35: 35, 36: 34, 37: 33},
    2: {31: 38, 32: 37, 33: 36, 35: 35, 36: 34, 37: 33},
    3: {31: 38, 32: 37, 33: 36, 35: 35, 36: 34, 37: 33},
    4: {31: 38, 32: 37, 33: 36, 35: 35, 36: 34, 37: 33}
}

# Process all predictions and store results
rmsd_results = []
final_rmsd_summary = {}

for pred_idx, pred_mol, mapping_pred in pred_mappings:
    active_overrides = manual_overrides_dict if APPLY_OVERRIDES else {}
    rmsd_before, rmsd_after = process_prediction_with_overrides(
        gt_mol, mapping_gt, pred_idx, pred_mol, mapping_pred, active_overrides
    )
    rmsd_results.append({
        "Prediction Index": pred_idx,
        "RMSD Before Override (A)": rmsd_before,
        "RMSD After Override (A)": rmsd_after
    })
    # Use the override result when enabled; otherwise this equals rmsd_before
    final_rmsd_summary[pred_idx] = rmsd_after

# ==============================================================================
# Step 6: Summarize Results and Create CSV
# ==============================================================================
print("--- Step 6: Generating Final Summary ---")

# --- Print table to console ---
if rmsd_results:
    print("\n--- RMSD Summary Table ---")
    print("-" * 70)
    print(f"| {'Prediction Index':<18} | {'RMSD Before Override (A)':<25} | {'RMSD After Override (A)':<25} |")
    print("-" * 70)
    for result in rmsd_results:
        idx = result['Prediction Index']
        before = f"{result['RMSD Before Override (A)']:.4f}"
        after = f"{result['RMSD After Override (A)']:.4f}"
        print(f"| {idx:<18} | {before:<25} | {after:<25} |")
    print("-" * 70)

# --- Print overall summary ---
if final_rmsd_summary:
    print(f"\n### Final RMSD Summary ({'after overrides' if APPLY_OVERRIDES else 'without overrides'}) ###")
    for pred_idx, rmsd in sorted(final_rmsd_summary.items()):
        print(f"Prediction {pred_idx}: RMSD = {rmsd:.4f} Å")
    
    best_pred_idx, best_rmsd = min(final_rmsd_summary.items(), key=lambda x: x[1])
    print(f"\nBest prediction: Index {best_pred_idx} with RMSD = {best_rmsd:.4f} Å")


# --- Save results to CSV file ---
csv_filename = 'rmsd_summary.csv'
headers = ["Prediction Index", "RMSD Before Override (A)", "RMSD After Override (A)"]

try:
    with open(csv_filename, 'w', newline='', encoding='utf-8') as csvfile:
        writer = csv.DictWriter(csvfile, fieldnames=headers)
        writer.writeheader()
        writer.writerows(rmsd_results)
    print(f"\n'{csv_filename}' has been saved successfully.")

except Exception as e:
    print(f"\nAn error occurred while saving the CSV file: {e}")

# ==============================================================================
# Step 7: Clean up generated files (Optional)
# ==============================================================================
# print("\n--- Step 7: Cleaning up generated PDB and coordinate files ---")
# files_to_delete = [
#     "gt_ligand.pdb", "pred.model_idx_0_ligand.pdb", "pred.model_idx_1_ligand.pdb",
#     "pred.model_idx_2_ligand.pdb", "pred.model_idx_3_ligand.pdb", "pred.model_idx_4_ligand.pdb"
# ]
# # Also add the coordinate files to the list
# for i in range(5):
#     files_to_delete.append(f"gt_coordinates_pred{i}.csv")
#     files_to_delete.append(f"pred{i}_coordinates_pred{i}.csv")
#     files_to_delete.append(f"gt_coordinates_pred{i}_override.csv")
#     files_to_delete.append(f"pred{i}_coordinates_pred{i}_override.csv")
#
# for file in files_to_delete:
#     if os.path.exists(file):
#         try:
#             os.remove(file)
#             print(f"✅ Deleted: {file}")
#         except OSError as e:
#             print(f"❌ Error deleting {file}: {e}")

print("\n--- Script Finished ---")
```
