## Supplementary_Data.zip: inputs/outputs, PyMOL sessions, and pocket-aligned RMSD workflow for "Single-sequence deep learning delivers crystal-quality models of covalent K-Ras G12 hotspot complexes": predictions_align_to_pocket.py

```
#!/usr/bin/env python

# -----------------------------------------------------------------------------
# Sungwon Jung
# -----------------------------------------------------------------------------

# This script run inside PyMOL to aligns multiple prediction objects to a reference pocket structure
# using only the alpha carbon (CA) atoms for the alignment.
# It aligns prediction objects named "pred.model_idx_0" through "pred.model_idx_4" to a pre-defined selection named "pocket".  

from pymol import cmd

def align_all_pred_to_pocket_ca():
    # Reference selection: pre-defined selection named "pocket" (CA atoms only)
    ref_sel = "polymer and name CA and (pocket)"
    
    # Iterate over pred.model_idx_0 through pred.model_idx_4
    for i in range(5):
        # Build the mobile selection for CA atoms in each prediction object
        pred_obj = f"pred.model_idx_{i}"
        mobile_sel = f"polymer and name CA and ({pred_obj})"
        # Name of the alignment object to store the result
        out_obj = f"aln_pred.model_idx_{i}_to_pocket"
        
        print(f"Aligning {pred_obj}: '{mobile_sel}' to '{ref_sel}' (object: {out_obj})")
        # Run alignment: align mobile_sel onto ref_sel and apply the transform to the whole pred object
        cmd.align(mobile_sel, ref_sel, quiet=0, object=out_obj, reset=1)

# Run alignment automatically on import
align_all_pred_to_pocket_ca()
```
