## Supplementary_Data.zip: inputs/outputs, PyMOL sessions, and pocket-aligned RMSD workflow for "Single-sequence deep learning delivers crystal-quality models of covalent K-Ras G12 hotspot complexes": README.md

Pocket-aligned RMSD (definition, workflow, and script roles)

Definition (as in AF3 SI): We define the binding pocket as all heavy-atom coordinates within 10 Å of any heavy atom of the ligand in the reference (ground-truth) structure. Predicted models are rigid-body–superposed to the reference using Cα atoms of residues within this pocket. RMSD is then computed over all heavy atoms of the ligand.

Reproducible workflow

1) Define the pocket in PyMOL (from the ground truth)
	1.	Load the ground-truth structure (e.g., PDB/CIF) and select the ligand (temporarily named sele by PyMOL).
	2.	Create the pocket selection (residue-expanded neighborhood around the ligand, excluding hydrogens):

select pocket, <GT_OBJECT> and (byres (sele around 10)) and not elem H

Replace <GT_OBJECT> with the ground-truth object name. Keep sele as the currently selected ligand.

2) Align predictions to the pocket (PyMOL)
	•	Load the five prediction objects named (you might want to change the name in case of using AF3 predictions):

pred.model_idx_0, pred.model_idx_1, pred.model_idx_2, pred.model_idx_3, pred.model_idx_4


	•	Run the alignment helper (aligns each prediction to pocket using only Cα atoms):

run path/to/predictions_align_to_pocket.py

After running, all five predictions are pocket-aligned to the ground truth.

3) Save ligands from GT and predictions (PyMOL)
	•	cd in PyMOL to the directory where you want outputs written.
	•	Edit variables inside save_ligands.py to match your data:
		•	gt_selection = "<GT_OBJECT> and resn <GT_LIG_RESN>"
		•	selection   = "<PRED_OBJECT> and resn <PRED_LIG_RESN>"
		•	Examples for <GT_LIG_RESN> include Y63, S08, or 1wu (use the actual residue name in your files).
	•	Run:

run path/to/save_ligands.py


	•	Expected outputs:
	•	gt_ligand.pdb extracted from the ground-truth object (e.g., “8T4V” or “7au1”).
	•	pred.model_idx_0_ligand.pdb … pred.model_idx_4_ligand.pdb extracted from each prediction.

4) Compute ligand RMSDs (Python)
	•	Use the script (RDKit + NumPy) to calculate RMSD between the reference ligand and each pocket-aligned predicted ligand. Optional manual overrides allow you to pin atom-to-atom correspondences when needed (e.g., different ordering, tautomers, altLocs). I used made this option to correct a mismatch from K-Ras(G12D) and G12Di.

python pocket-aligned_ligand_RMSD.py


	•	Outputs a CSV (e.g., pocket-aligned_ligand_RMSD.csv) 
	•	Sanity checks (bond counts, element sets, atom counts) are logged; any manual pairing overrides are recorded.

⸻

Script roles (what each file does)

	•	pocket-aligned_ligand_RMSD.py
Converts a Jupyter workflow into a script for batch RMSDs between a ground-truth ligand and multiple predicted poses. Uses RDKit (I/O, matching) and NumPy (RMSD). Reads PDBs, applies optional manual atom-pair overrides, performs checks, and writes a CSV summary.

	•	predictions_align_to_pocket.py (run inside PyMOL)
Aligns pred.model_idx_0 … pred.model_idx_4 to the pocket selection of the ground-truth object using only Cα atoms for the least-squares superposition.

	•	save_ligands.py (run inside PyMOL)
Saves ligands from the ground truth and from each prediction to separate PDBs. You edit gt_selection and selection to match your object names and ligand residue name (e.g., resn Y63 / S08 / 1wu). Produces gt_ligand.pdb and pred.model_idx_#_ligand.pdb.
