## Supplementary_Data.zip: inputs/outputs, PyMOL sessions, and pocket-aligned RMSD workflow for "Single-sequence deep learning delivers crystal-quality models of covalent K-Ras G12 hotspot complexes": save_ligands.py

```
#!/usr/bin/env python

# -----------------------------------------------------------------------------
# Sungwon Jung
# -----------------------------------------------------------------------------

# This script run inside PyMOL to save ligand structures from ground truth and prediction objects.
# It extracts the ground-truth ligand from object "7au1" with residue name S08, and save as "gt_ligand.pdb".
# It also extracts ligands from prediction objects "pred.model_idx_0" through "pred.model_idx_4" with residue name S08,saving them as "pred.model_idx_0_ligand.pdb" ... "pred.model_idx_4_ligand.pdb".


from pymol import cmd

def save_ligands():
    # Print currently loaded objects (for debugging)
    objs = cmd.get_names("all")
    print("Loaded objects:", objs)
    
    # Save Ground Truth ligand (object '7au1', residue name S08)
    gt_selection = "7au1 and resn s08"
    count_gt = cmd.count_atoms(gt_selection)
    print("Ground Truth selection ('7au1 and resn s08') atom count:", count_gt)
    if count_gt == 0:
        print("Warning: Ground Truth selection returned 0 atoms. Please check object '7au1' and residue name 'S08'.")
    gt_filename = "gt_ligand.pdb"
    cmd.save(gt_filename, gt_selection)
    print("Saved", gt_filename, "from selection:", gt_selection)
    
    # Save prediction ligands (residue name S08 in each object)
    for i in range(5):
        obj_name = f"pred.model_idx_{i}"
        selection = f"{obj_name} and resn s08"
        count_pred = cmd.count_atoms(selection)
        print(f"Selection count for {selection}:", count_pred)
        if count_pred == 0:
            print(f"Warning: selection '{selection}' returned 0 atoms. Check object '{obj_name}' and residue name 'S08'.")
        out_filename = f"{obj_name}_ligand.pdb"
        cmd.save(out_filename, selection)
        print("Saved", out_filename, "from selection:", selection)

# Run immediately upon load
save_ligands()
```
